# Energy efficient convolutional neural networks for arrhythmia detection

**DOI:** 10.1101/2021.09.23.461522

**Authors:** Nikoletta Katsaouni, Florian Aul, Lukas Krischker, Sascha Schmalhofer, Lars Hedrich, Marcel H. Schulz

## Abstract

Electrocardiograms (ECG) record the heart activity and are the most common and reliable method to detect cardiac arrhythmias, such as atrial fibrillation (AFib). Lately, many commercially available devices such as smartwatches are offering ECG monitoring. Therefore, there is increasing demand for designing deep learning models with the perspective to be physically implemented on these small portable devices with limited energy supply. In this paper, a workflow for the design of small, energy-efficient recurrent convolutional neural network (RCNN) architecture for AFib detection is proposed. However, the approach can be well generalized to every type of long time series. In contrast to previous studies, that demand thousands of additional network neurons and millions of extra model parameters, the logical steps for the generation of a CNN with only 114 trainable parameters are described. The model consists of a small segmented CNN in combination with an optimal energy classifier. The architectural decisions are made by using the energy consumption as a metric in an equally important way as the accuracy. The optimisation steps are focused on the software which can be embedded afterwards on a physical chip. Finally, a comparison with some previous relevant studies suggests that the widely used huge CNNs for similar tasks are mostly redundant and unessentially computationally expensive.

## 1. Introduction

Monitoring, analysis and classification of the heart electrical activity have attracted the interest of the scientific community and became a field with a variety of commercial applications [1, 2]. Small portable devices, such as smartwatches, or implantable heart recorders [3] are capable of monitoring a heart’s rhythm and activity. Their small size and their high production and placement costs require hardware with low energy consumption. Consequently, the embedded software on these devices, which is responsible for the detection of abnormal heart rhythm (arrhythmia), must have restricted computational requirements.

Atrial fibrillation (AFib) is a type of arrhythmia caused by disorganised atrial functionality. It is the most common cardiac arrhythmia with a rate of 1% in the general population [4]. AFib can be diagnosed by the electrocardiograph (ECG), as the irregular, fast heartbeat leads to the absence of the P-wave, irregularities of the R-peaks and quite often in narrow QRS complexes. Although these fibrillatory waves are one of the major causes of strokes, early diagnosis of AFib and prompt treatment can inhibit the risk adequately [5].

Artificial intelligence has permitted the design of models which are able to address this issue instantly, by classifying in real-time the ECGs, indicating different types of heart arrhythmia and giving recommendations for further investigation and treatment by a cardiologist. Before the extended usage of deep learning, researchers alluded models for the automatic detection of arrhythmia based on heavy feature extraction strategies, which are application specific and require domain knowledge. Nowadays, the Convolutional Neural Networks (CNNs) and the Recurrent Neural Networks (RNNs) are the most common approaches used for the detection of miscellaneous types of arrhythmias, with results that pledge high performances. The success of CNNs is mainly due to their ability to “learn” all the essential features and classify them accordingly.

Hannun et al. [6] proposed a 34-layer CNN for the detection and categorisation to rhythm classes with higher accuracy than trained cardiologists. Before this, other studies used CNNs to develop accurate models for arrhythmia classification and Afib detection [7, 8, 9, 10]. Furthermore, recurrent connections between the segments of ECG signals were used in [11] and [12]. In the former study the final dense layer, that is responsible for the decision, was replaced by a Support Vector Machine (SVM), while in the latter one additional attention layer was included. Finally, skip connections were used by Xiong et al. [13].

However none of the above mentioned studies, whose main objective is to detect accurately these heart abnormalities, has considered the resulting energy consumption of these models which is an important aspect once they are placed on portable, wearable devices. Specifically, when the only concern is the accuracy of the CNN it is straightforward that deeper architectures will perform better, given that more detailed features are detected (certainly if overfitting is avoided). But when we are interested in the implementation of the model on actual integrated circuits this is a point that we should contemplate. Having energy efficient hardware components can obviously minimise the need for energy supply but software-wise speaking implementing huge models with hundreds of thousands of neurons and millions of connections is almost certainly energy-inefficient.

Some previous studies, in different domains of application, have already addressed this issue by developing techniques that simplify the CNN architecture and decrease the number of weights. Structured sparsity is first mentioned at the early years of neural networks [14]. Since then, multiple works have used it to compress the network’s architecture. Han et al. [15] proposed a pruning strategy with quantisation of the trained weights in order to enable weight sharing and Huffman encoding. They achieved four times layerwise speedup and seven times more energy efficiency. A structured pruning method by particle filtering on kernels and feature maps was introduced by Anwar et al. [16]. The different structures were evaluated by the classification accuracy with proved good performance on small CNNs. The not important parameters were excluded in the studies of Alvarez et al. [17] and Zhou et al.[18] taking advantage also of the structured sparsity. Another approach for generating simplified versions of neural networks while maintaining all the predicting capabilities of the bigger ones is the Knowledge Distillation(KD). In the works of Bucila et al. [19] and [20] KD was applied and a smaller *student* network was trained simultaneously with a much bigger *teacher* network by optimising the loss function between them, proving the satisfactory performance of the *student* network although its much smaller, compressed architecture.

These methods propose a lightweight version of the initial network. Despite the benefits they may have by reducing the network size, they are mainly considering the efficiency of the network only after training. Supposing we care about a future implementation on hardware, we are interested on a stable architecture that can be updated by changing the model parameters, while maintaining the basic structure. By pruning the network weights and architecture in a second step, the generalisability of the network is affected and a future update will demand drastic intervention on the network design. And this is a condition that results to additional time and financial burdens for the hardware producers. Therefore, we deem it necessary to consider the energy efficiency while designing the models. An attempt in this direction was done by Amirshahi et al. in [21], where an ECG classification algorithm was developed for energy efficient wearable devices with the use of spiking neural networks. They suggested the transformation to the spike domain by encoding the heartbeat signals into spikes and using the spike-timing dependent plasticity to train the weights of the layers according to the spike timings. They show that, since the calculations are done in the spike domain, the energy consumption is significantly reduced.

In this study we are presenting an accurate energy-efficient architecture to detect arrhythmia with minimised number of nodes and connections in a way that a possible implementation of the model on physical chips can benefit low energy constraints and actual area. For the purpose of the paper we apply our method on the detection and classification of atrial fibrillation. The input ECG signals are segmented to non-labeled windows of equal length. Although the labels of our dataset are provided per signal, the networks are generated in such a way that can detect Afib per segment. The models can lead to devices with more durability, less charge cycles and reduced computation power while at the same time the high detection capabilities are preserved. Our method is not scenario specific. It can be applied on every kind of time series, generalised to more classes, different kinds of inputs and addresses the aspect of energy-efficient neural networks assessing that large architectures are mostly redundant. In the context of this paper, we are focusing on the software implementation with features that will allow us to embed it later on a physical chip. In the next sections the exact workflow for energy-optimised models is described. The performances are analysed and compared to other relevant recently published studies, which suggest networks with thousands of extra nodes and millions of additional trainable parameters.

## 2. Materials and method

### 2.1. Overview

In the following subsections, the workflow for the design of energy-efficient CNNs is presented. Our method suggests a pipeline for the construction of CNNs for time series that has as guideline not only the precision in the detection but also the energy efficiency. The energy consumption, defined here by the network size, number of computations and amount of trainable parameters, contributes to the choice of the final model architecture and it is coupled with the training of the network. The network optimisation consists of a grid search in thousands of models, network segmentation and application of the optimal energy classifier. An overview is shown in Figure 1. The exact steps are the following:

1. preprocessing of the input signal for noise reduction,
2. construction of multiple model architectures with a grid search for different number of filters, filter kernels, layers and pooling sizes.
3. training of the models
4. comparison of the candidate models using as a metric the accuracy and the energy consumption. The choice of the candidate models is done by setting an accuracy threshold and searching for the ones that minimise the energy consumption.
5. model segmentation to enable predictions per segment
6. find best parameter values for the optimal energy classifier

**Figure 1:**
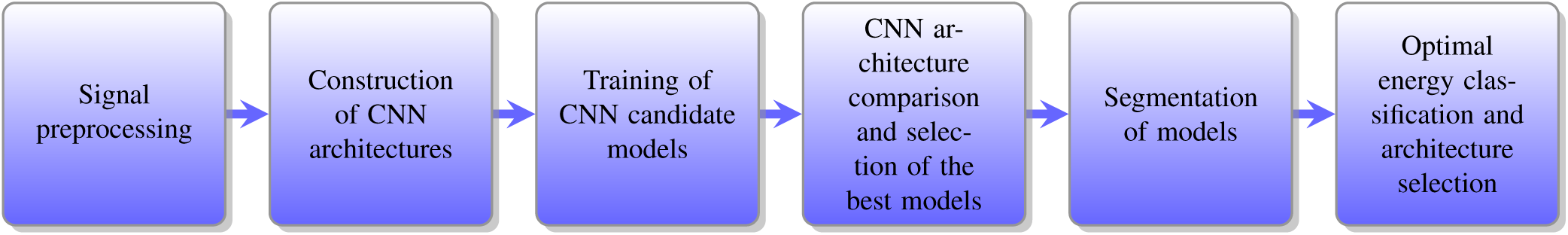
Workflow for the generation of energy-efficient models for time series.

After the last steps of energy optimisation (model segmentation, optimal energy classifier) the optimal model which has the fewest trainable parameters while preserving high accuracy can be selected.

### 2.2. ECG dataset

The data used for this study was provided by the Bundesministerium für Bildung und Forschung(BMBF)^1^ in the context of the project “Energieeffizientes KI-System”. The dataset consists of 16.000 ECG signals, 8.000 with AFib and 8.000 control cases of sinus rhythm. The signals were measured by a portable device PM1000, GETEMED AG and the two channel ECGs (leads I and II) were provided. Each of them has a duration of approximately 120 seconds with sampling rate 512 Hz. Examples of the signal data are shown in Fig. 2.

**Figure 2:**
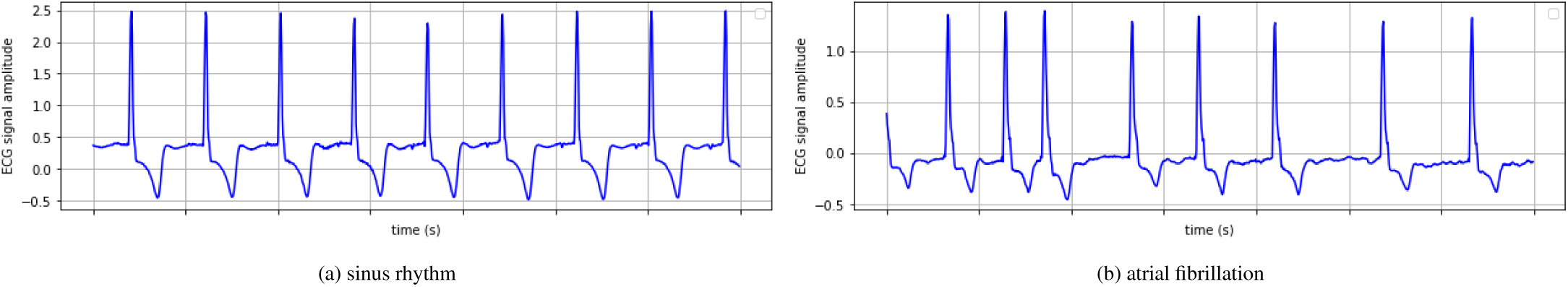
Two example ECG signals of the dataset, one of each class, are illustrated. The subfigure (a) is a control case that corresponds to the sinus rhythm and the subfigure (b) is an example of atrial fibrillation. In both of the cases one segment of 7 seconds is depicted.

### 2.3. Dataset with ECG signals

For the purpose of our approach the 2 minutes were sectioned into 17 segments with equal duration of 7 seconds. The 7 seconds duration for the windows was selected after trials, where we searched for the minimum necessary duration which can lead to accurate Afib detection. Though, the labels are assigned to the whole signal and not to each segment. The AFib signals have not only persistent fibrillatory waves in the entire duration but also paroxysmal events, where AFib can be detected only for some seconds and then the normal rhythm recurs. It is observed that at the beginning of the ECGs, a noisy, not periodical wave is appearing as a results of the placement and initial calibration of the device. Thus, the first segment of each of the ECGs was excluded by the training process. The rest of the noise that arises by the hardware measurement device or by the human movement was handled partially by the preprocessing step and by our proposed model that is robust to disturbed signals.

### 2.4. Band-pass filter for ECG signal preprocessing

One of the main difficulties that we should overcome when monitoring continuous ECG signals is the noise by muscle stimulators, magnetic fields, corrupted signal caused by electrode misplacement, baseline wander or even noise generated by the respiration of the individuals. In order to distinguish the main artifacts of the ECG from the noise a band-pass Butterworth filter was applied on the raw signal.

In order to enable the transfer of our architecture design on a physical hardware, the preprocessing strategy was chosen properly. A 14th order Infinite Impulse Response (IIR) filter was used, which could be energy efficiently implemented in analog or digital hardware. The parameters of the 7 biquad blocks of the second order stage (SOS) architecture can be determined to achieve the wanted transfer function of the bandpass filter. The high order of the filter enables a very good suppression of baseline errors and noise. The future hardware implementation will of course need a proper rearrangement of the second order stages to keep the amplitude in between the filter stages in a reasonable range.

The application of the filter was done on the frequency domain by eliminating all the frequencies smaller than 5 and bigger than 30 Hz. In figure 3 one ECG signal is plotted before and after the denoising by the bandpass filter. By doing so, the CNN is capable of finding optimal patterns in both of the classes and extract features that differentiate them. The bandpass filter has not only a smoothing effect on the signal but also centers the signal around zero.

**Figure 3:**
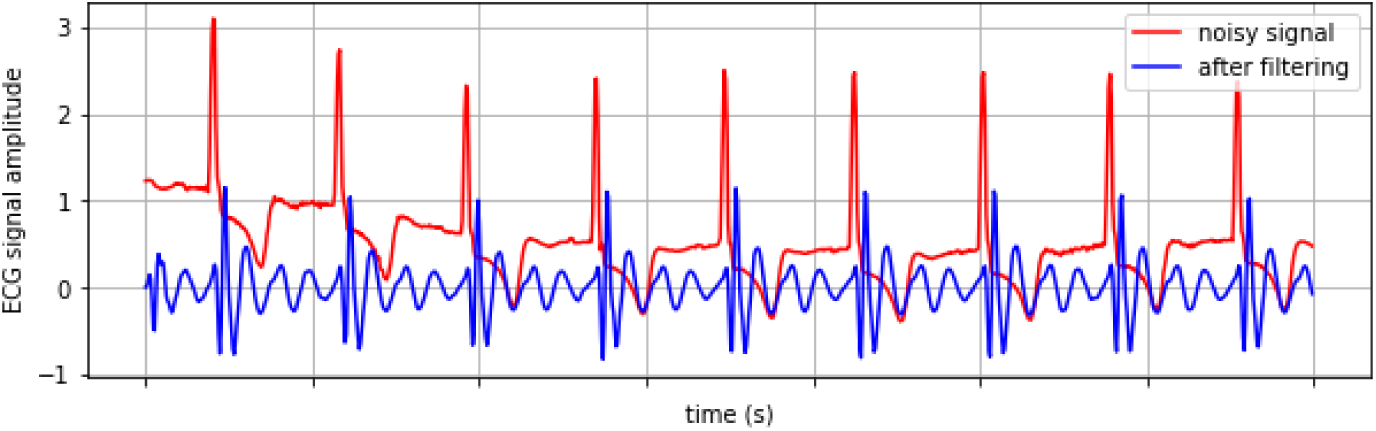
ECG signal before and after the application of the bandpass filter. The red line is the signal before the preprocessing. After the elimination of the frequencies smaller than 5 Hz and bigger than 30 Hz the “clean” signal is centered and shown in blue color.

### 2.5. Convolutional neural networks (CNNs)

CNNs are a popular type of deep learning models that have huge potential in a variety of disciplines. Many studies have used them for image and signal classification, object detection, signal denoising and many others. They are mainly consisting of three different operations constructed as layers. The *convolutional layer*, the *activation function* and the *pooling layer*. A CNN’s ability to extract highly complex and data-driven features for all the above mentioned scenarios is mainly due to the convolution operations. More precisely, each convolutional layer applies a cascade of filters, commonly known as kernels, on the input signal and is arranged in feature maps, each of which extracts different kinds of features.

Considering the complexity of the input signals, the linear nature of the convolution cannot capture all the underlying information. Therefore, the activation functions serve as a mapping of the previous layer to the next one in a non-linear manner. However, the application of multiple filters on the same input often dramatically increases the dimensions of the feature maps, thus the pooling operation is responsible of condensing the complexity of the CNN simply by down-sampling information. Commonly, the generated features of the CNN are fed into fully connected layers with dense connections between them. The number of the layers, the kernel, pooling size and the number of nodes in the fully connected layers are some of the hyperparameters, that define the structure of the CNN and should be chosen appropriately with regard to network performance and learning ability.

Nevertheless, when there is a need of energy efficiency and limited network size, it is recommended to minimise the number of nodes, edges and the overall computations, while preserving high accuracy. The property of CNNs to apply the same kernel along the whole signal without changing its weights is called *weight sharing*. This attribute can scale down excessively the number of trainable parameters that should be calculated during learning and also limit the memory requirements for the weight storage.

### 2.6. Initial CNN architecture

The first training of the candidate networks was done as follows. All the 16 segments of each signal are passed through the feature extractor of the CNNs. The output of the last pooling layer is flattened and fully connected to 4 nodes. At the end 4 ∗ 16 = 64 nodes were saved for the whole approximately 2 minutes signal and were fully connected to the output node using the sigmoid activation function. By doing so, we are forcing the CNN to learn general representations based on the whole signal, reducing the risk of filters which are performing well only locally (for example at the beginning or at the end of the signal).

### 2.7. CNN Learning

The initial CNNs were trained with the adaptive moment (Adam) optimizer [22]. For the convergence of the CNN to an optimal value, the binary cross entropy loss function was applied:

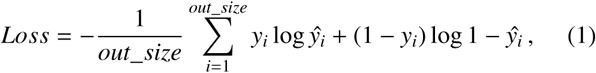

where *y* is the ground truth and *ŷ* the predicted class.

Also, a dropout layer with parameter 0.5 is implemented before the fully connected layer to avoid overfitting. This means that during the training only 50% of all the weights are updated at each iteration.

We used the following procedure to assess the performance of the models: The training set was randomly split into 80% for training, 20% for testing and 10% of the training subset was used for validation (see Tab. 1). Parameter optimization was done only on the validation set, whereas performance computation was done on the test set. By design it cannot happen that a full ECG, or a segment of it, from the same person is part of the training and test set. All ECGs which are part of the test or validation set, are independent of the ones used for training in order to avoid an overoptimistic performance assessment of the models.

### 2.8. CNN architecture comparison

Choosing the right CNN architecture can be quite challenging, because many hyperparameters need to be learned as mentioned above. In our application, the accuracy is not the only metric that we want to optimise. Although a model can be accurate enough for AFib detection, if its complexity is quite high and with many nodes and connections, then the energy consumption for the prediction of one ECG signal will be very large if the model will be integrated on a hardware with limited energy supply.

We conducted a grid search over number of filters, kernel size and pooling size to find the optimal architecture. By setting an accuracy threshold, we can pick models that are less complex but well performing. The complexity of the models is calculated in terms of neurons by the following equations:

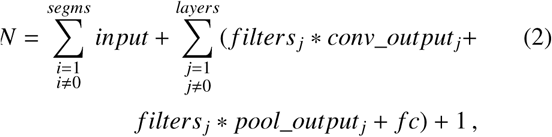

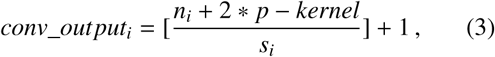

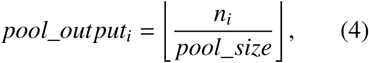

where *N* is the total amount of neurons of the CNN, *segms* denotes the number of segments for the signal, *layers* the number of layers (convolution, pooling), *filters* the number of filters, *pool* the pooling size and *fc* the nodes in the fully connected layer. The addition of 1 at the end of the equation corresponds to the output node. Concerning the calculation of the output for each convolution *conv_output* the *n* denotes the input to be convolved, *p* is the padding, *kernel* the size of the filters and *s* the stride for the application of the convolution. Finally the *pool_size* is the window size of the pooling operation.

By following this strategy, the CNNs with the higher accuracy and less complexity are selected. However, these are not the final architectures. Further optimisations are done in the next steps as described afterwards.

### 2.9. Recurrent CNN and energy optimisation

The model as it is described in the previous section requires the whole 2 minutes ECG signal in order to make the final decision. Howbeit, in cases of paroxysmal AFib the fibrillatory waves can be spotted only in some parts of the ECG. Assuming that the AFib is detected at the early seconds, it is straightforward that the process of the rest of the signal is meaningless and it should be classified as AFib. Feeding the whole ECG in the CNN needs extra computation power that in some cases is unnecessary. With regard to this fact, we developed a fully segmented model which is capable of making a decision for each of the 7sec windows (Figure 5). The new architecture is combining the output of the current segment with information by the previous segments to make decisions. Using segments in such a recurrent manner allows us to have a temporal dynamic behavior. At the end, although each segment is treated individually by the recurrent CNN (RCNN), the decision is made based on all the previous observations.

**Figure 4:**
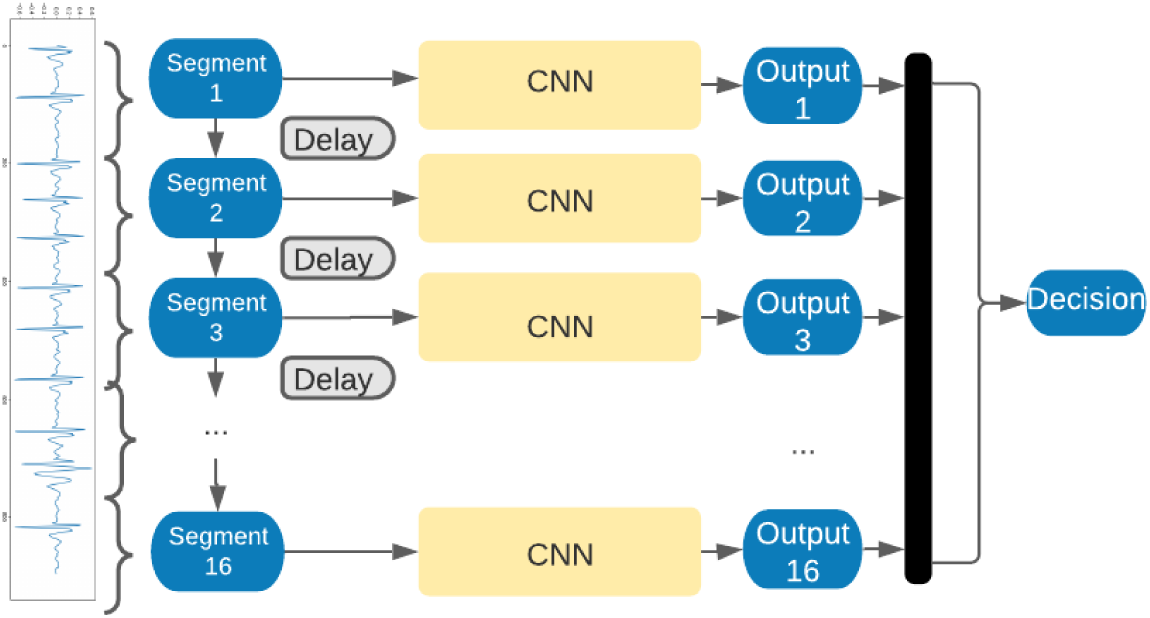
Model for segmented signal classification. In this architecture the input signal is segmented into windows of predefined length. The whole 2 minutes signal is fed into the same CNN in segments. For each segment a number of nodes is stored. Once all the segments are passed and their outputs are concatenated, the final classification is performed. As we do not have labels for each of the segments, this architecture allows us to train the network using only one label per signal while training the same network to extract the important features, using the information of all the segments.

**Figure 5:**
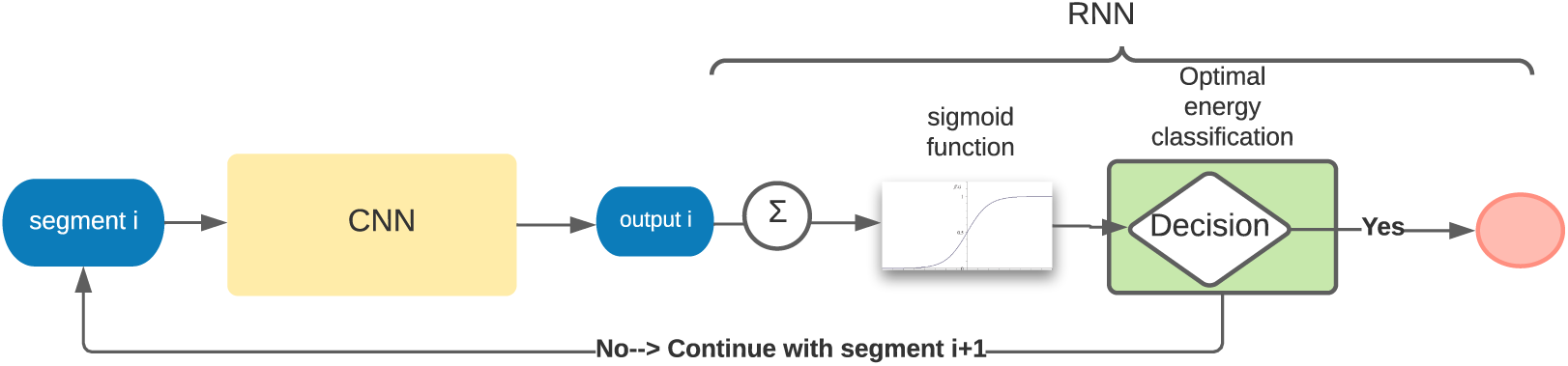
Recurrent CNN architecture with energy optimisation. The decision of our model is now done per signal segment, by considering recurrently the outputs of the previous segments. This recurrent flow of information is achieved by using the optimal energy classifier and transforms the network to a recurrent CNN. The recurrent module of the CNN has parameters that need to be optimised, as it is described in the Optimal energy classification subsection.

In more details, we are using transfer learning to train the RCNN. The nodes with their weights of the feature extractor part of the model are preserved and the part of the models until the last pooling is frozen. The flattened output is now fully connected to one output with the use of the sigmoid activation function. The output of the sigmoid though, is not the prediction of the model. The final decision is made by using the optimal energy classification approach (Figure 5), which is the memory and decision unit of our RCNN.

### 2.10. Optimal energy classification

In order to build a simplified small RCNN, we used the optimal energy classifier. The memory and decision unit of the RCNN has three additional parameters that need to be optimised. The detailed algorithm, presented in pseudocode 1, takes as input the *down limit D, upper limit U* and the *number of successive segments S* of the same class that should be detected in order to have a decision. These three values are set by a grid search on the training set. For example, let us assume that these three values are set to 0.47, 0.53, and 5. If the output of each segment after the application of the activation function is smaller than 0.47 the whole signal is classified immediately as sinus rhythm and no more segments are streamed into the network. If it is bigger than 0.53 the whole signal is classified as arrhythmia. In the case that the output of the current segment is between these two values, then the next 7 seconds segment is fed into the network until we get an output smaller than 0.47, bigger than 0.53 or 5 successive segments of possible arrhythmia (> 0.50) or 5 successive segments of possible no arrhythmia (< 0.50) (Alg. 1).

By doing so, we are permitting the RCNN to make a decision faster and shut it down. Even though our hybrid model does not contain any kind of complex modules for recurrent connection between the segments, like long short-term memory (LSTM) [23], we have managed to have a temporal dynamic behavior.

## Results

In order to test our method, a dataset of 16.000 ECGs measured by a portable device was used. The dataset was balanced for both cases and controls randomly split for training and testing. The ECG signals were divided into 80% for training, 20% for testing. Also, a 10% subset of the training set was used for validation. A more detailed description of the numbers can be seen in Table 1. The noise generated by the device, the movement of the patients and the respiration was eliminated by the application of a band-pass filter in the range of 5-30 Hz. The lengths of the signals were approximately 2 minutes with a sampling frequency of 512 Hz. After downsampling, the remained signals have a sampling rate of 128 Hz. The ECGs were segmented in 7 seconds windows. In total 16 segments per signal were acquired.

**Table 1:** Split of raw ECG signals.

|  | Training | Validation | Testing |
| --- | --- | --- | --- |
| Atrial fibrillation | 4994 | 554 | 2452 |
| Sinus rhythm | 5086 | 566 | 2348 |
| Total | 10080 | 1120 | 4800 |

### Algorithm 1 Optimal energy classification

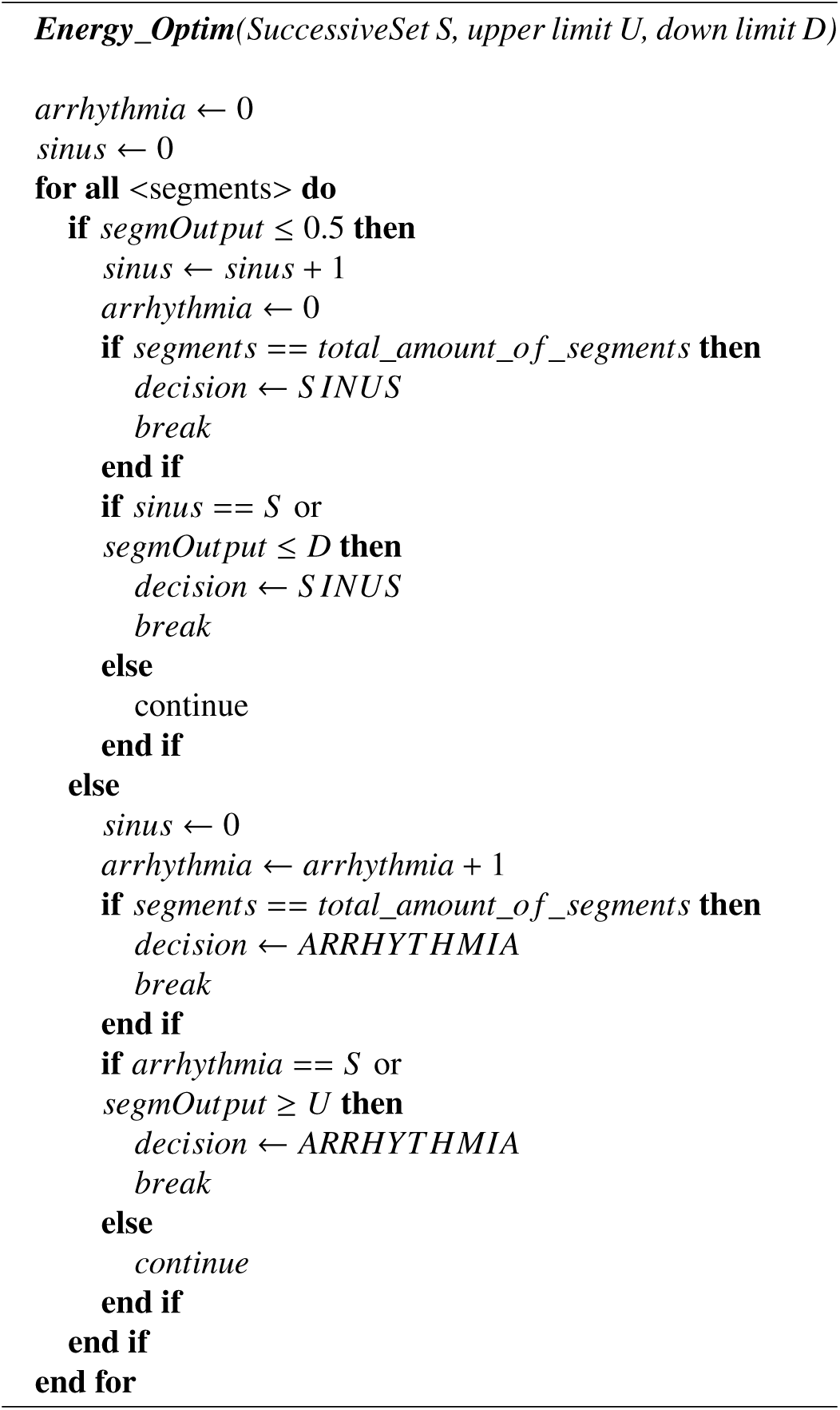

After denoising and normalising the signals, a grid search of multiple CNN architectures is performed to compare their performance. As the strategy is to keep the models small enough to allow low energy needs, we restricted the search on 3 layers. However the number of filters varied in the range of 1-5 for each of the layers, different kernel sizes between 4 and 11 were tested and pooling sizes between 2 and 6 were examined.

Due to absence of labels for each individual segment, the decision for each of the signal was made after passing all the 16 segments into the network. More precisely, each of the 16 segments was fed into the network successively. The output of the last convolutional-pooling layer is flattened, fully connected and stored to a predefined amount of nodes. These nodes of the fully connected layer were restricted in the range 2 to 5 for the grid search. These nodes are concatenated for all the segments and fully connected to the output (Fig. 4). In this way we allow the feature extraction to be learned on features by the whole signal and capture the important ones. In Fig. 6 a comparison between model accuracy and model complexity is depicted. The complexity of the models denotes the sum of a model’s nodes, see equation (2).

**Figure 6:**
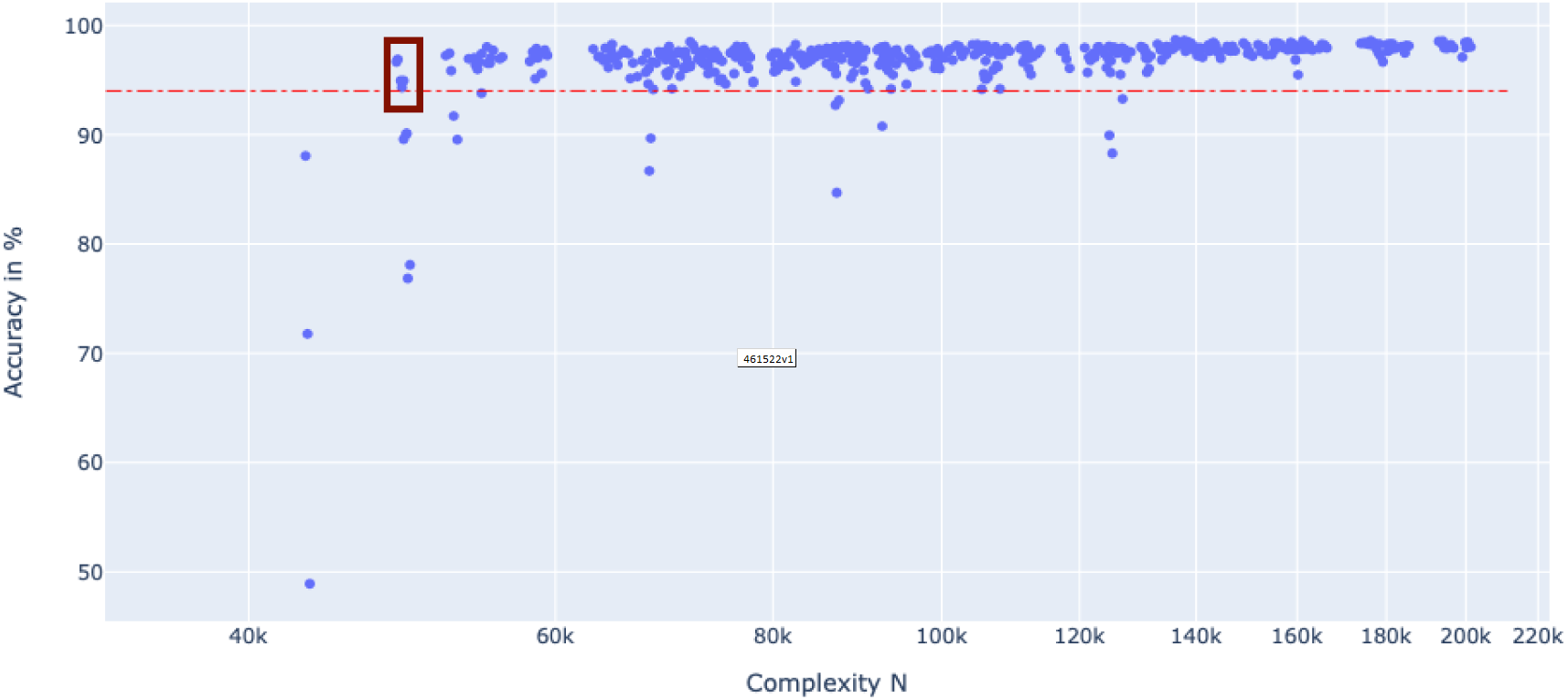
CNN model architecture comparison. A subset of the tested models is displayed. Each dot in the scatter plot is one distinct model with its architecture. By setting a threshold of 94% for the accuracy, the model with the smallest complexity above this threshold is considered to be the most energy-efficient. The red dashed line represents this threshold and the selected dots are the chosen candidate models that fulfill the requirements

Setting a threshold of 94% for the test set, which renders applicability in practice, the models with the least complexity are chosen. As depicted in Fig. 6, there are five models that have the fewest number of nodes, while preserving the accuracy above the threshold. These models are the candidates, which are selected to undergo the further energy optimisation steps when included into the RCNN, such that classification can be done potentially without reading all segments.

A more detailed description of the chosen models’ architectures can be found in Table 2. Specifically, all the models have three layers of convolutions with 1, 2 and 2 filters respectively. Each convolutional layer is followed by a ReLU activation function and a pooling layer for dimensionality reduction of size 3, 6 and 6. However, the kernel sizes of the convolutions are varying in the range of 7 to 11. The sizes of the filters are affecting the number of trainable parameters and computations in the network. Namely, by increasing the kernel size, an increased number of neighbour nodes will contribute to the current calculation.

**Table 2:** Network architectures of 5 most energy-efficient candidate models. All the models consist of 3 1D convolutional layers followed by pooling operations. The number of filters, the kernel and pooling sizes are displayed in the table. All the convolutional layers have ReLU as activation function and the dense layer a sigmoid.

| Model | Parameters |  |  |  |  |  |  |  |  |  |
| --- | --- | --- | --- | --- | --- | --- | --- | --- | --- | --- |
|  | Layer 1 |  |  | Layer 2 |  |  | Layer 3 |  |  | Dense layer |
|  | Conv1D filters | Conv1D kernel | Pooling size | Conv1D filters | Conv1D kernel | Pooling size | Conv1D filters | Conv1D kernel | Pooling size | Nodes |
| 1 | 1 | 8 | 3 | 2 | 8 | 3 | 2 | 8 | 6 | 1 |
| 2 | 1 | 9 | 3 | 2 | 9 | 3 | 2 | 9 | 6 | 1 |
| 3 | 1 | 10 | 3 | 2 | 10 | 3 | 2 | 10 | 6 | 1 |
| 4 | 1 | 11 | 3 | 2 | 11 | 3 | 2 | 11 | 6 | 1 |
| 5 | 1 | 7 | 3 | 2 | 7 | 3 | 2 | 7 | 6 | 1 |

## 2.11. Energy optimization for reduced energy consumption

Using one of the CNN architectures for the classification of arrhythmia in a continuous fashion, for example 12 hours while wearing a smart watch, would be very energy inefficient. In practice, it makes sense to limit the detection of arrhythmia to short repeated intervals, here we are using 7 seconds intervals, but this may differ. Ideally, the classifier can decide about arrhythmia or non-arrhythmia, without exploring the whole 2 minutes.

As the CNN’s feature extraction part has been trained on the whole 2 minutes signal, it has the ability to extract the necessary features for the detection of Afib. Having created an accurate classifier we need to force the model to make decisions independently per segment. Since the features of all the segments are extracted in the same way, we assume that if the flattened output of the last convolution-pooling is fully connected to one node then the weight of this node will reveal the decision of the segment. Therefore, by freezing the weights of all the convolutional layers (feature extractor), flatten the last convolution-pooling layer’s output, connect it to only one node (output-decision) and retrain only the last classification layer, we have a fully segmented model. The restriction of the absence of labels per segments can now be overcome by using the addition of all the segments’ outputs as a final decision (after the application of the sigmoid to restrict the values in the range of [0-1].

The integration of the optimal energy classifier into the CNN permits the judgement for each signal without processing all the 2 minutes. Using the signal segments in a recurrent way we can make decisions per signal window while considering at the same time the previous signal segments. The final RCNN architecture that is the combination of the previously described CNN and the optimal energy classifier is generated as following: For the training set the outputs after application of sigmoid for all the segments are saved. For multiple combinations of upper limit, down limit and number of successive segments (see Algorithm 1) the average number of needed segments per signal and the accuracy of all the candidates are computed. If the output of the segment is smaller than the down limit, or bigger than the upper limit the decision is immediately defined as Afib and no arrhythmia, respectively. Otherwise, the rest segments of possible Afib or no arrhythmia are needed. After testing all the combinations for the algorithm, we choose the best performing. For our models of interest the chosen parameters can be found in Table 3. The tested values for the down limit were set in the range 0.20 till 0.48 with step size 0.02 and for the upper limit in the range 0.52 till 0.80 with the same step size. The number of tested succesive segments was set in the interval of 2 till 8. One of the combinations that minimises the number of needed segments while retaining high accuracy is selected. The decision for the final model is done after all the optimisation steps are completed.

**Table 3:** Performance and size of the most energy-efficient RCNN models before and after optimisation. The accuracies for all the models before and after energy optimisation are presented for the training and test set. Also, the average number of 7 seconds segments that are needed for the whole 2 minutes ECG classification is provided. The average number of segments in this table corresponds to the results on the test set.

| Model | Performance before energy optimisation |  | Performance after segmentation and energy optimisation |  |  |  |  |  |  |  |
| --- | --- | --- | --- | --- | --- | --- | --- | --- | --- | --- |
|  | Training set | Test set | Training set | Test set | Down limit | Upper limit | Successive number of segments | Average number of segments | Energy reduction | Parameters |
| 1 | 0.951 | 0.943 | 0.926 | 0.918 | 0.30 | 0.80 | 4 | 4.798 | 70% | 93 |
| 2 | 0.952 | 0.949 | 0.914 | 0.920 | 0.40 | 0.60 | 6 | 3.844 | 76% | 100 |
| 3 | 0.974 | 0.968 | 0.934 | 0.946 | 0.40 | 0.60 | 6 | 3.276 | 80% | 107 |
| 4 | 0.974 | 0.966 | 0.956 | 0.953 | 0.40 | 0.60 | 6 | 3.092 | 81% | 114 |
| 5 | 0.960 | 0.949 | 0.936 | 0.942 | 0.42 | 0.58 | 4 | 5.702 | 64% | 88 |

The average number of segments, presented in Table 3, corresponds to the average number of 7 seconds segments needed by each of the models for the correct classification of the whole approximately 2 minutes ECG signals. Although the parameters of the algorithm for the optimal energy classification are calculated for each of the models on the training set, the average number of needed segments is estimated on the test set for an unbiased evaluation.

The most energy efficient RCNN was model 4 that has overall 114 distinct variable parameters (weights, biases and 3 parameters for the classifier), resulting to a very small architecture with the potential to classify long ECG signals. Also, in cases of continuous monitoring and classification, without shutting down the whole system after the decision, it can be a powerful, lightweight model for uninterrupted AFib detection. Using this architecture and the optimal energy classifier in the model, the RCNN’s classification decision on the test set was made only by streaming 3.092 segments on average. Instead of feeding the whole 2 minutes ECG in the model we can get an accurate prediction only by testing 27.44 sec. This can reduce the computational cost to almost 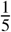 while preserving 95.3% accuracy.

Before the energy optimisation step, the model 5 seems to be the more efficient option, as it has a smaller kernel and consequently less operations are needed at each application of the filters. Howbeit, after the conversion to RCNN, with the use of the optimal energy classifier we need on average 5.702 segments for the classification, in contrast to model 4 that needs only 3.092. Hence, the use of 26 extra trainable parameters is an acceptable trade-off while considering the reduced number of segments needed.

The proposed RCNN architecture consists of 3 layers of convolutions with 1,2 and 2 filters followed by average pooling of size 3,3 and 6 and one output node with sigmoid activation function. The upper and down limits of the optimal energy classification are set to 0.60 and 0.40 and the parameter for the successive segments is chosen to be 4. It should be mentioned that the selection of these parameters is not absolute but they must be adjusted according to the predefined accuracy and energy restrictions for each application. Yet, it was a proper decision for our case study.

## 2.12. Comparison with recently published architectures

The main advantage of the proposed models through our workflow is their small size, not only in terms of their ability to classify long time series, but also in comparison with other recently published architectures in the same field of application. Chaur et al. [24] generated similarly an 1D CNN for the detection of atrial fibrillation. The CNN architecture consists of 10 layers of convolutions followed by pooling operations and 2 fully connected layers followed by one softmax layer output. The number of filters at each convolutional layer varies in the range of 32 to 512, which results in 3,933,634 trainable parameters. This denotes 34,505 times more parameters than our proposed model 4.

For a fair comparison with our approach, we reproduced the network architecture and trained it on our dataset with the suggested parameters. The network was trained using the Adam optimisation algorithm and cross-entropy as loss function. The batch size was fixed to 50 and the network was trained for 100 epochs. Due to absence of labels for each of the segments we cannot perform segment-wise training and therefore the full length signal is inserted as input. Anyhow this is not affecting the final number of parameters and network size, as all the segments had to be fed. We calculated the test set accuracy for comparison, where Chaur’s model achieved an accuracy of 98.8 %. This performance is 3,5 % higher than our model’s accuracy. Though, when the energy efficiency is equally important as an accurate detection rate, our model overpowers Chaur’s approach by a factor of 34,505 when it comes to trainable parameters. Considering these results, one must decide if the 3,5 % is a reasonable compromise for such a huge energy saving.

Additionally, we tried to generate the 1D-CNN as it is described in [25] and [26] for atrial fibrillation and trained it on our data. However, it was not feasible to achieve model convergence and produce a stable accurate solution. Their model comprises a total of 232,214,329 parameters, 13 layers of convolutions 2 fully connected and one sigmoid output layer. As this model has 2,036,967 more trainable parameters than our model, it is likely that the amount of training data was not sufficient for a model of this complexity.

Let us assume now that we want to generate a model with accuracy as high as Chaur’s model. In that case, a model with higher classification performance from Figure 6 can be selected. For this purpose we chose the model with 6, 6 and 7 filters, kernel size of 9 and pooling sizes equal to 3, 3 and 6 and named it as model 6. After the energy optimisation steps, this model has 98.2% on our test set and on average 2.47 segments are needed for the classification of one ECG signal. The final model has a total of 2,347 parameters. Although the accuracy is almost similar to Chaur’s proposed architecture, they used 3,931,287 more trainable parameters. This suggests that Chaur’s model architecture is highly redundant at least for the variations observed in our data. The architectures for comparison of Chaur’s model, our most energy efficient Model 4, and our Model 6 with the highest performance can be found in Table 4 and Figure 7.

**Table 4:** Architectures and performances of Chaur’s model and our proposed energy efficient Model 4 and the best performing Model 6.

| Model | Layers | Filters | Parameters | Accuracy |
| --- | --- | --- | --- | --- |
| energy-efficient model | 3 | 5 | 114 | 95.2% |
| best performing model | 3 | 19 | 2,347 | 98.2% |
| Chaur et al.[24] | 10 | 1,984 | 3,933,634 | 98.8% |

**Figure 7:**
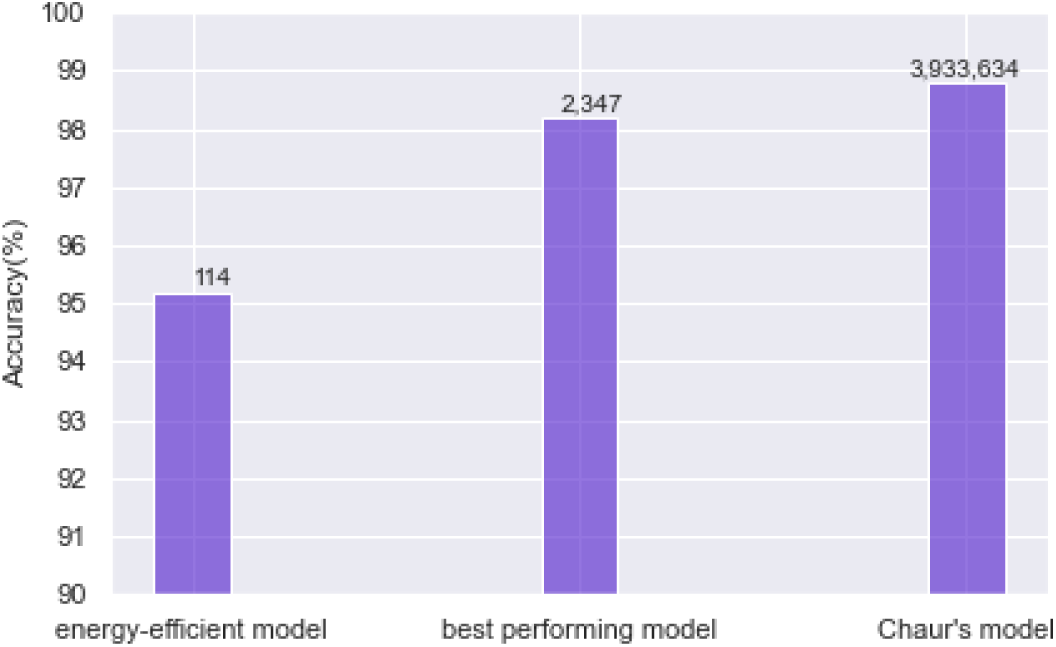
Comparison of the most energy efficient model using the proposed workflow (Model 4), the best performing model (Model 6) and Chaur’s model [24]. The numbers on top of the bars indicate the size of the model with respect to the trainable parameters.

## 3. Conclusion and Discussion

In the present study, we are proposing energy efficient recurrent CNN architectures for long time series and our approach is tested on the detection of atrial fibrillation on ECG signals. Our workflow suggests the development of lightweight, fully-segmented models with drastically fewer model parameters than previous studies. The inclusion of the energy consumption as an additional metric for the evaluation of the performance, allows us to generate architectures that can be easily embedded on physical small hardware devices.

Developing light-weight neural networks that can be incorporated on tiny chips and placed on wearable devices is a challenge, as we want to keep restricted energy requirements and high performance. In our method, the choice of the model architecture is not absolute. It can be done, by taking into consideration the accuracy and energy restrictions. In other words, it is a trade-off between accuracy and energy consumption. One should define these limitations beforehand. Afterwards, the architecture that better meets the current needs is selected.

The choice of the preprocessing method varies per task and nature of signals. For the current task of Afib detection, bandpass filtering was applied in the range of 5-30 Hz for signal denoising and normalization. The filtering step could be replaced by some extra layers of convolutions, but as the main idea of our implementation is to maintain a small network with as few neurons and parameters as possible, the band-pass filter was essential for noise canceling. While some previous works apply in a similar manner filtering of small and high frequencies [26], the transformation to the frequency domain by a Fourier or wavelet transform is also used [10]. With the intention of incorporating our designed model on a physical chip, the band-pass filter can offer an “inexpensive” solution, given that it can be applied directly in time domain, avoiding this way extra transformations and it is a well established method for analog and digital chips. Furthermore, a variety of papers are focusing on the detection of R-peaks [7]. We considered this approach as energy inefficient, in a way as the detection of spikes, demands many additional computations and it is difficult to be generalised to general time series.

The optimal architecture for our application consists of 3 convolutional layers with 1, 2 and 2 filters respectively and 1 fully connected layer. The total number of parameters of the model is 114, which is millions of times smaller than model sizes that others have suggested. After energy optimisation our model achieved an accuracy of 95.3% on our test set of 4800 ECGs. The use of the optimal energy classifier permitted us to reduce the energy by 81% for the classification of 2 minute signals. Specifically, only an average of 3.09 signal segments of 7 seconds, or approximately 21 seconds, were needed for the classification of the whole 112 seconds. Mistakes due to wrong segment-wise decisions are avoided by recurrently using the information of previous segments. Also, another architecture with The focus of the paper is on the model generation in means of software. It is describing a succession of steps that need to be followed, in order to facilitate the future mitigation of the model on a chip. As future work, we are concentrated on the transfer of the model on a simulated chip. This of course requires some additional optimizations such as weight quantization to fit the requirements of the chip technology, quantization of the filter coefficients to avoid numerical instabilities and an iterative approach for the correction of inaccuracies between the software and hardware implementation.

## Funding

We acknowledge funding from the Alfons und Gertrud Kassel-Stiftung as part of the center for data science and AI. This work was supported by the DFG Cluster of Excellence Cardio Pulmonary Institute (CPI) [EXC 2026].

## Footnotes

1 https://www.bmbf.de/

## Notes

### Competing Interest Statement

The authors have declared no competing interest.

